# HIV-1 antisense protein of different clades induces autophagy and associates to the autophagy factor p62

**DOI:** 10.1101/320341

**Authors:** Zhenlong Liu, Cynthia Torresilla, Yong Xiao, Clément Caté, Karina Barbosa, Éric Rassart, Shan Cen, Benoit Barbeau

## Abstract

Over recent years, strong support argues for the existence of an HIV-1 protein encoded by antisense transcripts and termed Antisense Protein (ASP). Furthermore, a recent *in silico* analysis has provided evidence for its recent appearance in the genome of HIV-1. We have previously detected ASP in various mammalian cell lines by Western blot (WB), flow cytometry and confocal microscopy analyses and reported that it induced autophagy, potentially through multimer formation. The aim of the current study was to examine autophagy induction by testing ASP from different clades, and to identify the implicated autophagy factors. We firstly confirmed that NL4.3-derived ASP was interacting with itself and that multimer formation was dependent on its amino region. Removal of this region was associated with reduced level of induced autophagy, as assessed by autophagosome formation but deletion of the most amino cysteine triplet did not totally abrogate multimer and autophagosome formation. Expression vectors of ASP from different clades were next tested and led to detection of monomers and varying levels of multimers with concomitant induced autophagy, as determined by increased LC3-II and decreased p62 (SQSTM1) levels. Through confocal microscopy, ASP was noted to co-localize with p62 and LC3-II in autophagosome-like cellular structures. CRISPR-based knock-out of ATG5, ATG7 and p62 genes led to increased stability in the levels of ASP. Furthermore, co-immunoprecipitation experiments demonstrated the interaction between p62 and ASP, which was dependent on the PB1 domain of p62. Interestingly, immunoprecipitation experiments further supported that ASP is ubiquitinated and that ubiquitination was also responsible for the modulation of its stability. We are thus suggesting that ASP induces autophagy through p62 interaction and that its abundance is controlled by autophagy- and Ubiquitin/Proteasome System (UPS)-mediated degradation in which ubiquitin is playing an important role. Understanding the mechanisms underlying the degradation of ASP is essential to better assess its function.

**Author Summary:** In the present study, we provide the first evidence that a new HIV-1 protein termed ASP when derived from different clades act similarly in inducing autophagy, an important cellular process implicated in the degradation of excess or defective material. We have gained further knowledge on the mechanism mediate the activation of autophagy and have identified an important interacting partner. Our studies have important ramification in the understanding of viral replication and the pathogenesis associated with HIV-1 in infected individuals. Indeed, autophagy is implicated in antigen presentation during immune response and could thus be rendered inefficient in infected cells, such as dendritic cells. Furthermore, a possible link with HIV-1-associated Neurological Disorder (HAND) might also be a possible association with the capacity of ASP to induce autophagy. Our studies are thus important and demonstrate the importance in conducting further studies on this protein, as it could represent a new interesting target for antiretroviral therapies and vaccine design.

## Introduction

The genome of HIV-1 expresses essential genes for its replication, which include gag, pol and env, common to all replication-competent retroviruses in addition to the genes encoding regulatory proteins (Tat and Rev) crucial for virus replication. Furthermore, HIV-1 is known to encode a number of accessory proteins, i.e. Vpu, Vpr, Vif, and Nef, all of which are suggested to be implicated in the inactivation of restriction factors and the modulation of immune functions (1–6). A former study had however demonstrated that HIV-1 isolates contained a conserved open reading frame (ORF) on the antisense strand of the Env gene, thereby hinting toward the existence of a tenth gene with a encoding potential for a 189 amino acid protein (7, 8). A recent study has further underscored the high degree of conservation of this ORF in most HIV-1 clades, but further argued that this gene correlated with the spread of the virus, being less present (or absent) in most SIV, HIV-1 groups N, O and P and less prevalent HIV-1 clades (9). Importantly, a number of previous and more recent studies have further confirmed that antisense transcription overlapping this ORF sequence was detected in transfected and chronically infected cells (10–19).

The existence of the presumed encoded protein termed ASP (<u>A</u>nti<u>s</u>ense <u>P</u>rotein) in infected patients has been further suggested by several groups through detection of specific antibodies and CD8+ T cell-mediated immune responses (10, 20–22). However, ASP detection itself has proven to be more difficult and was first analyzed by in *vitro* translation and electron microscopy in transfected and chronically infected cells (10, 15). More recent studies have provided a better understanding of this protein in terms of its potential membrane localization and its biased expression in monocyte-derived macrophages and dendritic cells (7, 17). In another recent report, we have been able to clearly detect the ASP protein for the first time by Western blot and various other approaches (confocal microscopy and flow cytometry) in transfected cell lines (23). Furthermore, our work further suggested that ASP was unstable, formed multimers and induced autophagy, likely through the formation of these high molecular weight complexes.

Autophagy, a major cellular degradation pathway, plays an important role in developmental processes, cellular stress responses, and immune pathways induced by pathogens. The early step of what is known as macroautophagy initiates through mTOR inhibition and ensuing formation of an active autophagy complex consisting of the phosphorylated Beclin-1 factor, normally negatively inhibited by its binding to Bcl-2 (24–27). Following this activation, a protrusion of two lipid-based structures termed the isolation membrane and the omegasome is induced intracellularly to form the initial opened structure termed the phagophore (28). The phagophore further elongates circularly and finalizes the maturation process of the autophagosome, in which are trapped cytosolic elements targeted for degradation (27). Final fusion of the autophagosome with lysosomes leads to degradation of its content and recycling of amino acids and other constituents (29, 30). Different proteins play an active role in autophagy, such as phagophore-associated Autophagy-related-genes (ATG) ATG5, ATG7, ATG10 ATG12 and ATG16 (31). One of the classical autophagy marker is LC3-II, a phosphatidylethanolamine-modified form of LC3-I form, which is embedded in the autophagosome membrane and contribute to its formation. Another implicated autophagy marker is p62/SQSTM1, a scaffold protein, which links aggregated complexes to LC3 and consequently becomes itself degraded.

Autophagy can either be beneficial or detrimental toward viral replication; the outcome of induced autophagy depends on the virus, the cell type, and the cellular environment (32). In fact, for HIV-1, initial studies in CD4+ T cells had shown that the envelope gp41 subunit induced autophagy-dependent apoptosis of uninfected cells (33, 34). However, a genomic screen has also identified certain autophagy-related host factors, which were rather essential for HIV-1 infection (35). Furthermore, in macrophages specifically, HIV-1-induced autophagy greatly improved gag pr55 processing and particle production (36). Importantly, Nef has been reported to block the process of autophagy at late stages (fusion with lysosomes) and avoid intracellular degradation of viral particles (5, 37). More recent data has also demonstrated a potential role for other HIV-1 proteins in regulating autophagy, such as Vif (38, 39).

Although previously associated to the proteasome degradation pathway, ubiquitination, the covalent conjugation of ubiquitin to proteins, has become of high relevance to autophagy (40–43). The C-terminus of ubiquitin is covalently link to the target protein by specific lysine residues (e.g. K48, K63), which subsequently leads to polyubiquitination. While K48- and K29-linked polyubiquitination has been shown to be optimal for degradation through the proteasome, other types of lysine-linked polyubiquitinations (e.g. on K63, K11, K6) and monoubiquitination may regulate processes such as autophagy, translation and DNA repair (43). In fact, ubiquitinated proteins can be recognized by the autophagy pathway through specialized adaptor proteins, also called sequestosome-1 like receptors (including p62/SQSTM1), Optineurin and NDP52, among others). The p62 protein can multimerize via its PB1 domain (NH_2_-terminal Phox and Bem1p domain) and bind LC3-II via its LC3-interacting region (LIR) and ubiquitinated proteins through the phosphorylation of its ubiquitin-associated domain (UBA) (42–44).

Based on our previous report linking ASP to autophagy, in the current study, we aimed at examining the mechanism behind ASP-induced autophagy. Our results demonstrate that multimerization of ASP and induced autophagy partly involves its amino end. We show that ASP from various HIV-1 clades all induced autophagy, as determined by LC3-II levels and p62 degradation. We further highlight that ASP co-localized and interacted with p62 through its PB1 domain and finally present data supporting that ASP is ubiquitinated, further contributing to its targeting toward degradation pathways.

## Results

**Analyses of ASP multimers**. A limited number of studies have shown that ASP could be detected in infected and transfected cells (15, 23). Using a codon-optimized ASP expression vector, we have successfully detected the ASP protein in transfected mammalian cells by several technical approaches, and concomitantly, demonstrated that it was forming multimers and could induce the formation of autophagosomes, thereby providing an explanation for its difficult detection (23). Herein, we sought to further define the mechanism leading to autophagy induction by ASP and to extend these analyses to ASP from different HIV-1 clades.

We were first interested in confirming the self-multimerization potential of ASP using expression vectors for Myc-tagged and chimeric GFP-optimized ASP (Fig. 1). Extracts from co-transfected 293T cells were immunoprecipitated with anti-Myc antibodies and subsequently analysed in parallel with total extracts by Western blot. As depicted in Fig. 1A, a specific signal was detected corresponding to the GFP-tagged ASP in immunoprecipitated extracts, which could not be accounted by significant difference in expression of the fusion protein in the different analysed samples. Similar co-transfection experiments were conducted in COS-7 cells and again revealed that both ASP proteins were associated (Fig. 1B). As cysteine-rich region of the protein might mediate these high molecular complexes, we pretreated the extract from Myc-tagged-ASP expressing COS-7 cells with high concentrations of reducing agents, DTT and β-mercaptoethanol. Upon treatment, the high molecular signals were strongly diminished with either agent and resulted in increased intensity of the 20 and 200kDs signals. Very high concentration of DTT led to disappearance of the high molecular weight signals (Fig. 1C)

**Figure 1:**
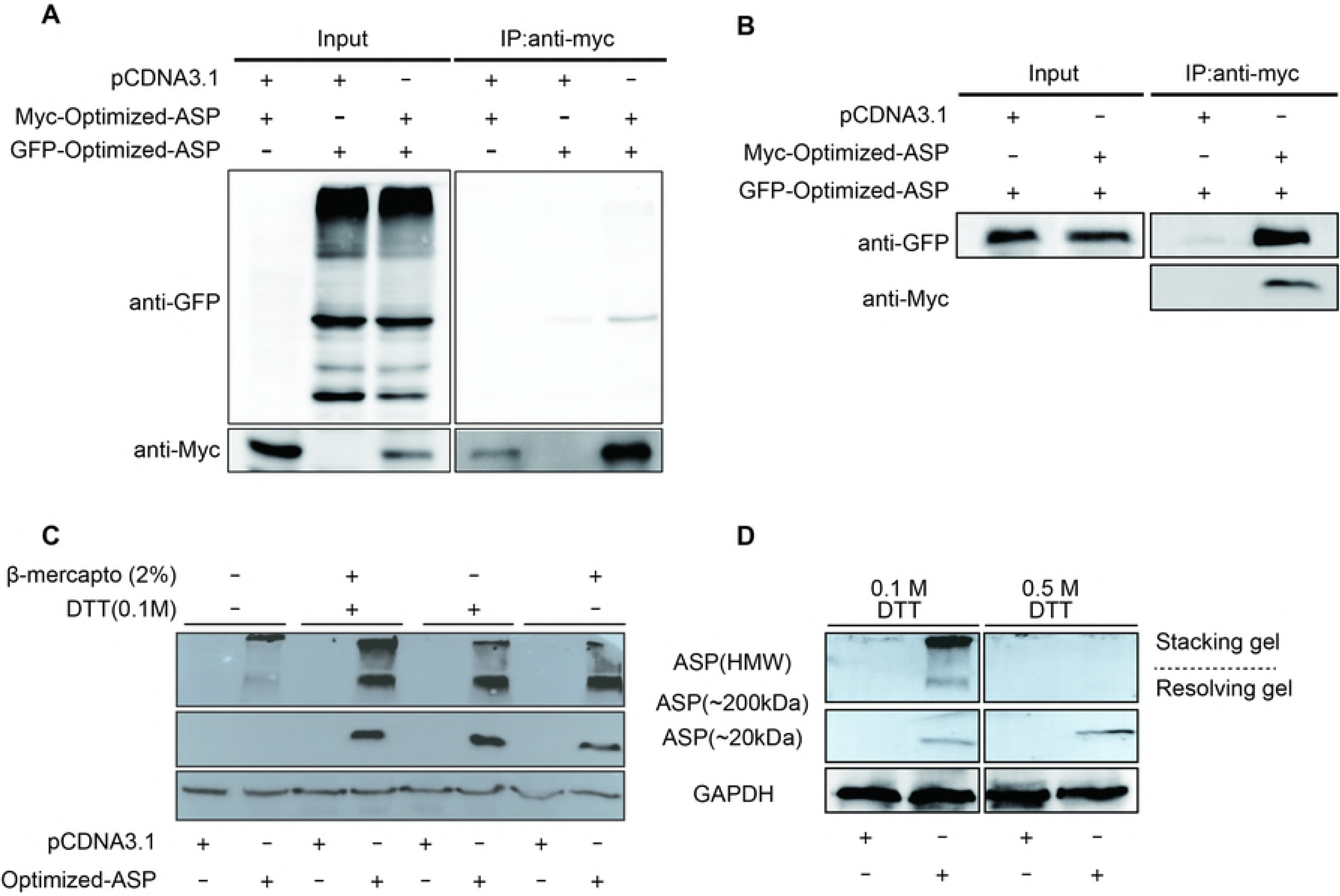
Multimerization of ASP involves disulfide bonds. **A-B**. 293T (A) and COS-7 (B) cells were transfected with expression vectors for GFP-optimized-ASP, Myc-tagged-optimized-ASP and/or the empty vector pcDNA3.1. At 48 h post transfection, immunoprecipitation was performed using the anti-Myc antibody and Western blot analyses were conducted through anti-GFP and anti-Myc antibodies. Total cellular extracts were similarly analysed in parallel with an anti-GFP antibody. **C**. Extracts from COS-7 cells transfected with the Myc-tagged optimized ASP expression vector (vs. pcDNA3.1) were treated with of DTT and/or β-mercaptoethanol prior to Western blot analyses with anti-Myc and anti-GAPDH antibodies. Both stacking and resolving gels are depicted and high molecular weight (including 200 kDa multimers) vs. monomeric 20kDa ASP signals are indicated.

In light of these results, a series of deletion mutants were next generated, in which the first 15 and 30 amino acid residues from the amino or carboxyl end of Myc-tagged ASP were deleted. Importantly, pMyc-optimized-ASPΔN1-15 and pMyc-optimized-ASPΔN1-30 vectors permitted the expression of an ASP mutant, from which 4 and 7 conserved cysteine residues were removed, respectively (Fig. 2A). Upon transfection of COS-7 cells, flow cytometry analyses showed that mutants and wild-type ASP were detected at comparative levels, except for mutant pMyc-optimized-ASPΔN1-30, which showed a reduced signal (Fig. 2B). The multimerization potential of these mutants were next analysed by Western blot (Fig. 2C). Interestingly, the loss of the first 15 residues had an important effect on the presence of high multimers, and led to a modest increase in the abundance of the monomer, while no such effect was observed with mutants bearing deletion at their COOH end. The ASPΔN1-30 mutant presented reduced signal for both monomeric and high molecular weight signals when compared to wild-type-expressing cells, which was likely due to reduced overall abundance of the protein. Confocal microscopy analyses were next performed in transfected cells (Fig. 2D). Deletion of the first 15 amino acids reduced (but not completely abolished) the presence of punctuated ASP signals, previously having been identified as autophagosomes. Deletion of the carboxyl end did not lead to differences in the distribution of the resulting ASP mutant from the wild-type version (Fig. 2D). Another mutant in which the conserved PXXP domain (position 47–53) was also tested in transfected 293T cells and revealed no clear change in multimer formation or induction of autophagy (data not shown).

**Figure 2:**
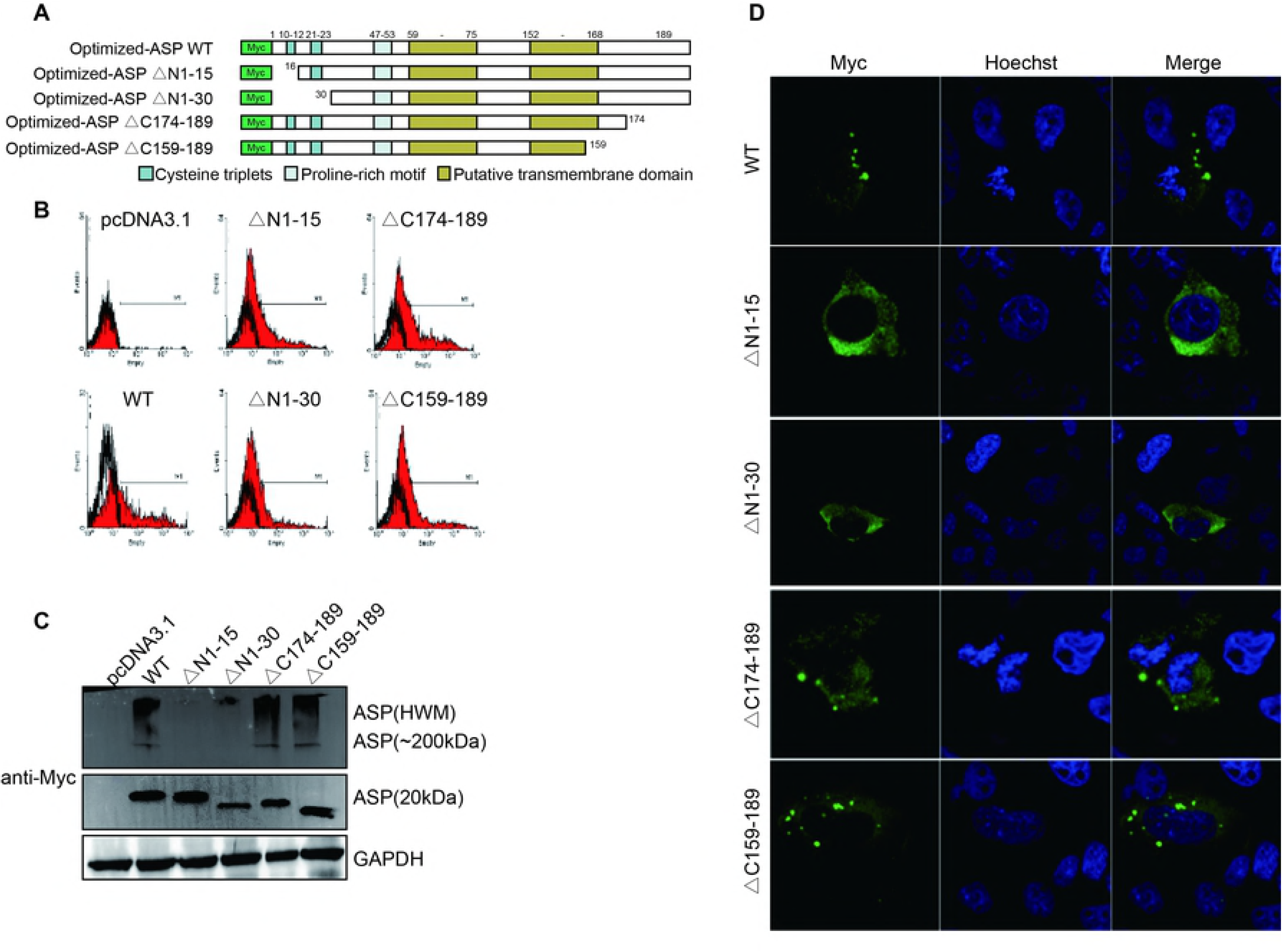
The amino end of ASP is implicated in its multimerization. **A**. Schematic representation of the different domains of ASP and the generated deletion mutants targeting the first 15 or 30 amino acids of either the N-or C-terminal of ASP. **B-D**. COS-7 cells were transfected with expression vectors for these different ASP mutants and the wild-type version vs. pcDNA3.1 (empty vector) and, at 48 h post-transfection, were analysed with an anti-Myc antibody by flow cytometry (**B**) and confocal microscopy (60X objective and with numerical aperture of 1.4) (**C**). Cells analysed by confocal microscopy were also stained for their nuclei with Hoechst. Cellular extracts from these transfected cells were also analysed by Western Blot using anti-Myc and anti-GAPDH antibodies (**D**). High molecular weight (including 200 kDa multimers) vs. monomeric 20kDa ASP signals are indicated.

Since a cysteine triplet is present in the first 15 amino acid region, we specifically deleted these cysteine residues (Fig. 3A). Flow cytometry indicated that the resulting mutant was detected at levels equivalent to those of optimized-ASP WT in transfected COS-7 cells (data not shown). Expression of the optimized ASP ΔCCC mutant in COS-7 cells resulted in a typical punctuated distribution in the cytoplasm (Fig. 3C), while Western blot analyses demonstrated that multimerization capacity of ASP ΔCCC appeared less pronounced with a concomitant increase in the monomeric signal, suggesting that this cysteine triplet was contributing to ASP multimer formation but that other cysteine (and possibly non-cysteine residues) were implicated in ASP multimerization and autophagy induction (Fig. 3B). We further tested the amino deletion mutants and the cysteine deletion mutants in 293T cells and confirmed in a clearer manner that the amino end and cysteine residues were affecting ASP multimer formation and protein levels (Fig. S1).

**Figure 3:**
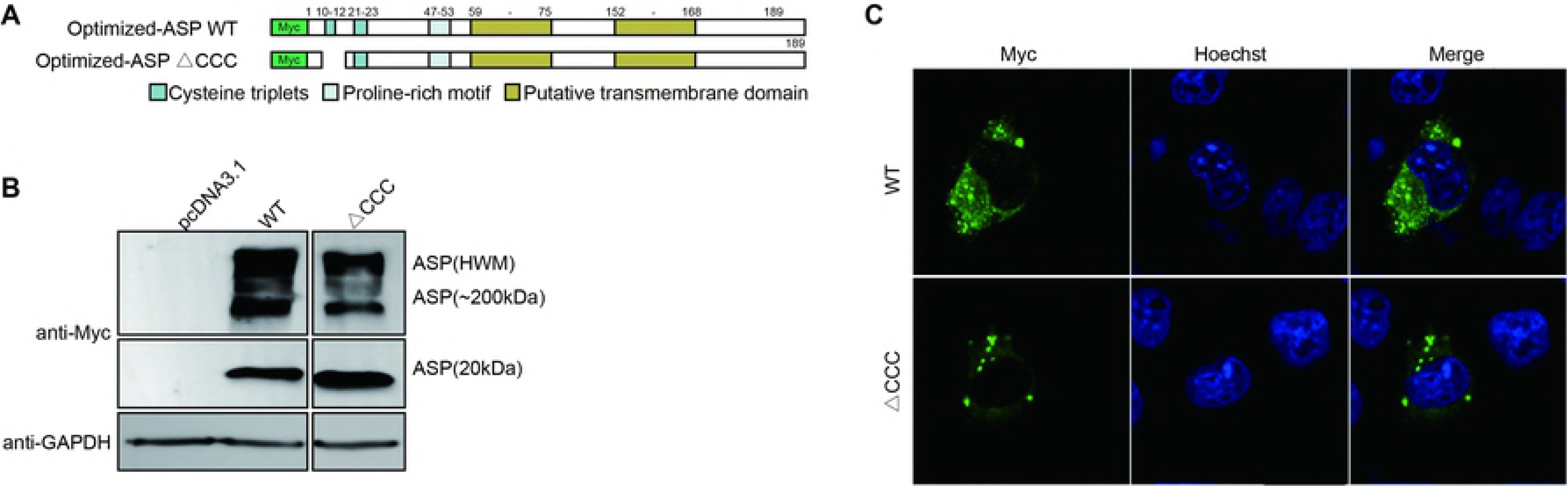
Deletion of the first cysteine triplet of ASP reduces multimer formation and autophagosome signals. **A**. Schematic representation of the different domains of ASP and the deletion mutant of the first cysteine triplet (^10^CCC^12^). **B-C**. COS-7 cells were transfected with expression vectors for wild-type or cysteine triplet-deleted ASP vs. pcDNA3.1 (empty vector). After 48h of transfection, using anti-Myc antibodies, cellular extracts were analysed by Western Blot (**B**) and confocal microscopy (60X objective and with numerical aperture of 1.4) (**C**). Cells analysed by confocal microscopy were also stained for their nuclei with Hoechst. High molecular weight (including 200 kDa multimers) vs. monomeric 20kDa ASP signals are indicated.

In light of these results, we conclude that the amino terminus (and cysteines) of ASP contribute to its capacity to multimerize, and is partly implicated in its autophagy-inducing properties.

**Expression and detection of ASP from various clades**. In order to better assess and understand the mechanism leading to ASP-mediated autophagy in a more representative manner, we next generated expression vectors for His-tagged ASP representing different HIV-1 clades. Importantly, these ASP ORFs were directly derived from the original proviral DNA sequence and were not codon-optimized, as we had previously performed in our earlier study (23). Sequence comparison of the ASP genes tested in this study is presented in Fig. 4A and highlights the previously observed absence of the first 25 amino acid in clade A ASP (9). This thereby allowed us to assess the multimer formation and the autophagy-inducing capacity of an ASP version naturally missing the typical amino end. Following transfection in 293T cells, we first used our monoclonal anti-ASP antibody derived against the epitope indicated in Fig. 4A to determine if ASP could be detected by Western blot. As depicted in Fig. 4B, despite variation in the amino acid sequence of the epitope in between tested ASP, we demonstrated the presence of the expected 20 kDa signal (albeit with some variation in molecular weight and intensity between clades). Clade A ASP was detected at a lower molecular weight signal in comparison to other clade ASP, except for the 92NG083 ASP. This signal was not due to preferential usage of the internal methionine residue, as demonstrated by the similar monomeric signals observed for all tested ASP upon analyses with an anti-His antibody (Fig. 4C) but could be accounted by lower amino acid number of this ASP in the variable COOH end region. In addition, high molecular weight signals including important aggregates were detected in cells transfected with the different ASP expression vectors. Interestingly, clade A showed a more abundant monomeric form when compared to the intensity of the multimers versus ASP from other clades. A similar behavior was noted for the clade A/G ASP representative (92NG083).

**Figure 4.**
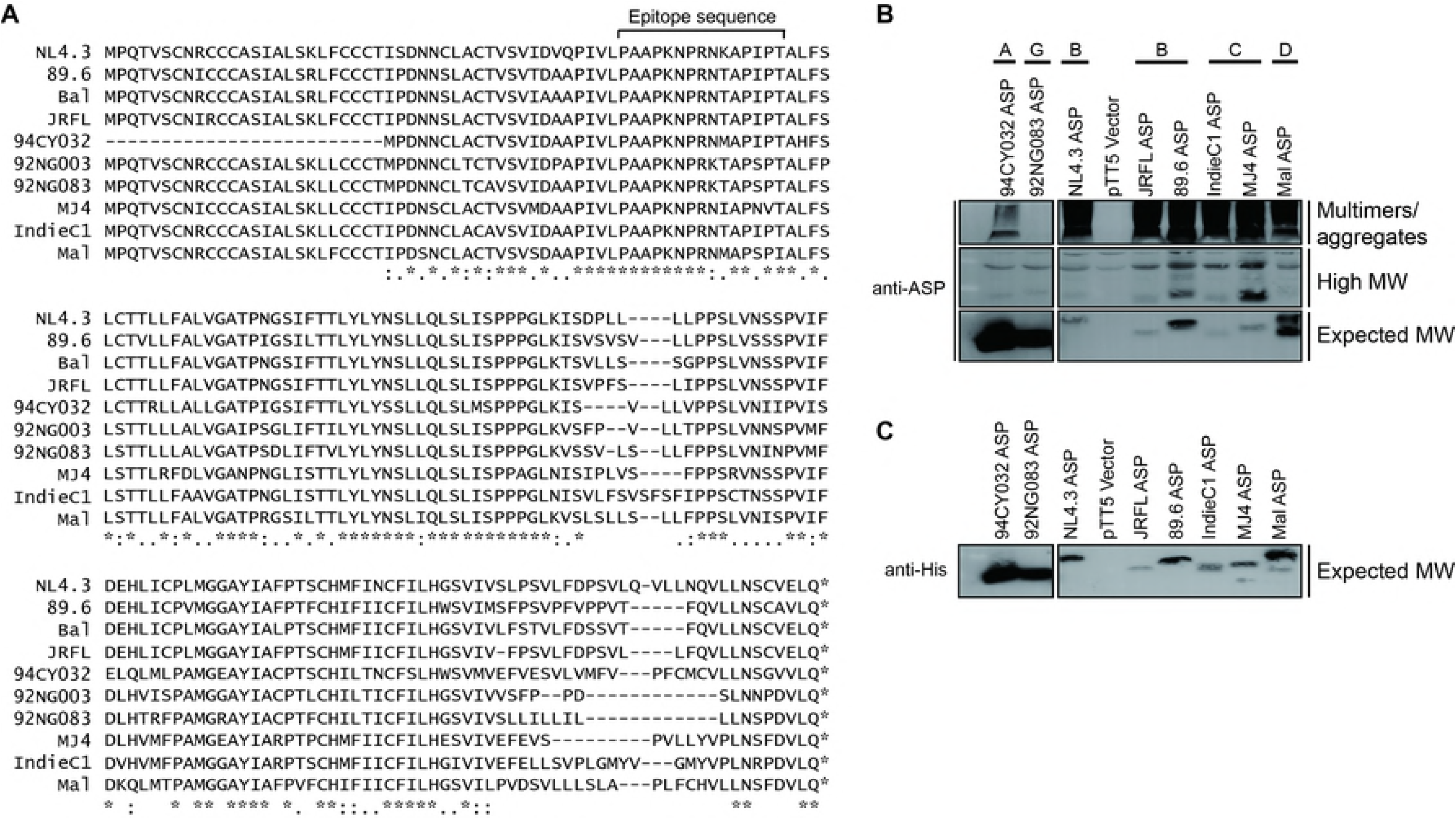
ASP from different clade representatives form detectable multimers. **A.** Sequence alignment of the predicted amino acid of ASP from different HIV-1 isolates representing different clades. The peptide region used to generate anti-ASP monoclonal antibodies is highlighted over the sequence from amino acid 47 to 61. **B-C.** Expression vectors of His-tagged ASP from various HIV clades were transfected in 293T cells and cell lysates were subsequently analyzed by Western blot for ASP (**B**: anti-ASP and **C**: anti-His). Depicted on the right side of the panels are multimers and high molecular weight and monomeric 20kDa ASP signals. Clade type for each tested ASP is indicated on top of each lane.

These results hence showed that His-tagged ASP from different clades can be detected with anti-ASP and anti-His antibodies. Furthermore, these results confirmed that multimer formation likely involved the amino end, but that, based on the variation of multimer intensities among various tested ASP (including results with the clade A/G representative), other regions of ASP are affecting the extent of multimer formation.

**Induction of autophagy by ASP from different HIV-1 clades**. Since the expression of all tested ASP could be detected and led to variation in the extent of multimer formation, we were next interested in determining if these different ASP proteins could indeed induce autophagy. Based on our previous results showing that levels of the lipid-modified microtubule-associated protein 1 light chain 3 (LC3-II), a well-known autophagy marker, was increased in 293T and COS-7 cells expressing ASP (23), we used this same marker to evaluate the autophagy-inducing capacity of ASP from different clade representatives. As depicted in Fig. 5A, the presence of ASP was again confirmed for all transfected ASP vectors and was detected as monomers and multimers. Importantly, ASP from the different clades all increased levels of LC3-II, albeit to different extent in transfected 293T cells. Furthermore, based on these Western blot analyses, clade A ASP 94CY032 seemed equally capable of inducing autophagy, when compared to other ASP expression vector. We then sought to validate that this increase in LC3-II levels was related to autophagy induction and not inhibition of the late step of the autophagy, both of which would lead to increase in the abundance of LC3-II levels. Hence, transfected 293T cell were treated with Bafilomycin A1, a blocking agent of lysosomal degradation. In these conditions, ASP-dependent induced levels of LC3-II should be maintained if ASP led to induced autophagy, while no such increase should be noted if ASP inhibited the last stage of autophagy. In Figure 5B, Western blot analyses revealed that LC3-II levels were indeed increased upon Bafilomycin A1 treatment, as expected but that ASP-driven induction was also evidenced. LC3-I/LC3-II ratios in ASP-expressing cells were calculated and shown to be higher than for similarity treated cells (−/+ Bafilomycin A1) transfected with the empty vector.

**Figure 5.**
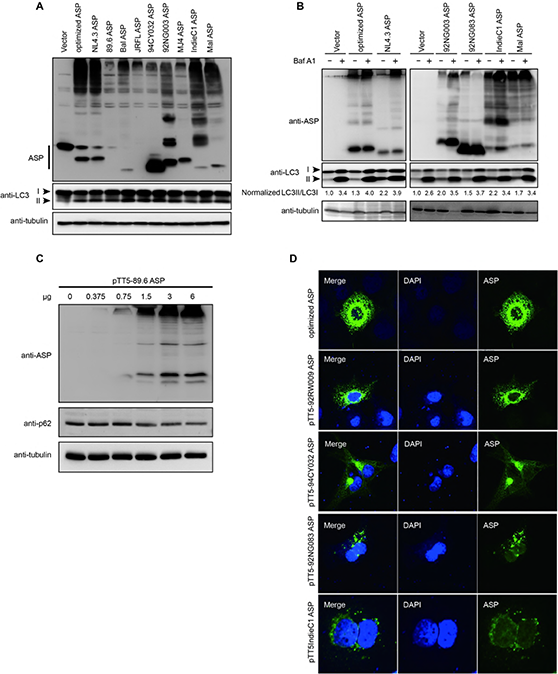
ASP from different clade representatives induce autophagy. **A-B.** Expression vectors of His-tagged ASP from various HIV clades and the empty vector were transfected in 293T cells and cell lysates were analyzed by Western blot for the detection of ASP (anti-His), LC3-II levels (anti-LC3) and tubulin. Signals for LC3-I, LC3-II and ASP are indicated on the left of each panel. In panel B, after transfection, cells were treated with DMSO (-) or with 100 nM Bafilomycin A1 (+) (Baf A1) for 6 h. LC3-II/LC3-I ratios are indicated for each lane and are representative of fold value over the calculated ratio for untreated cells transfected with the empty vector. **C.** Increasing quantity of clade 89.6 ASP expression vector was transfected in 293T cells and cell lysates were analyzed by Western blot for ASP, p62 and tubulin. **D.** COS-7 cells were transfected with expression vectors of Myc-tagged ASP from various clades. After fixation, cells were labeled with anti-Myc antibodies followed by goat anti-mouse IgG coupled to Alexa Fluor 488, stained with DAPI and observed by confocal microscopy.

Since p62 degradation can also be used as a marker of autophagy, we conducted additional Western blot analyses in 293T cells transfected with increasing quantity of the clade B Bal ASP expression vector. A dose-dependent decrease in p62 levels was confirmed upon ASP expression, further confirming induced autophagy (Fig. 5C). Expression vectors of ASP from different clades further revealed reduced levels of p62 in transfected cells when compared to empty vector-transfected cells (Fig. S2). As a further indication of autophagy being induced by these different ASP-expressing vectors, transfected COS-7 cells were analyzed by confocal microscopy using anti-Myc antibodies. Analyses revealed the characteristic presence of cytoplasmic punctuated ASP (including cells expressing clade A ASP), which is reminiscent of autophagosomes (Fig. 5D).

These results confirmed that ASP from different clades induced autophagy and that no striking differences were noted in between clades, especially with respect to clade A ASP. They further confirmed that ASP induced autophagy, and do not suggest an inhibitory action of ASP on late stage o autophagy leading to degradation of autophagosome content.

**Knockout of autophagy factors increase the abundance of ASP**. Using inhibitors of early and late steps of autophagy, we have previously demonstrated that the abundance of codon-optimized NL4.3 ASP increased (23). In order to further confirm the involvement of autophagy in the regulation of ASP, we first conducted targeted knock-out of the ATG5 and ATG7 genes using the CRISPR/Cas9 system by lentivirus-mediated transduction (Fig. 6). 293T cells were first stably transduced with lentivirus expressing selective sgRNA and following puromycin selection, two clones per knock-out were subsequently transfected with the His-tagged NL4.3 ASP expressing vector. Resulting extracts were analyzed for ASP, ATG5 and ATG7 expression. Knock-out of ATG5 or ATG7 through targeting of two different regions resulted in the reduced levels of the ATG5-ATG12 complex, although more limited for clone sgATG5-1 and, of ATG7, respectively (Fig. 6A and B). Importantly, His-tagged ASP levels increased in these clones, with a more increased abundance associated with lower expression levels of the targeted gene.

**Figure 6.**
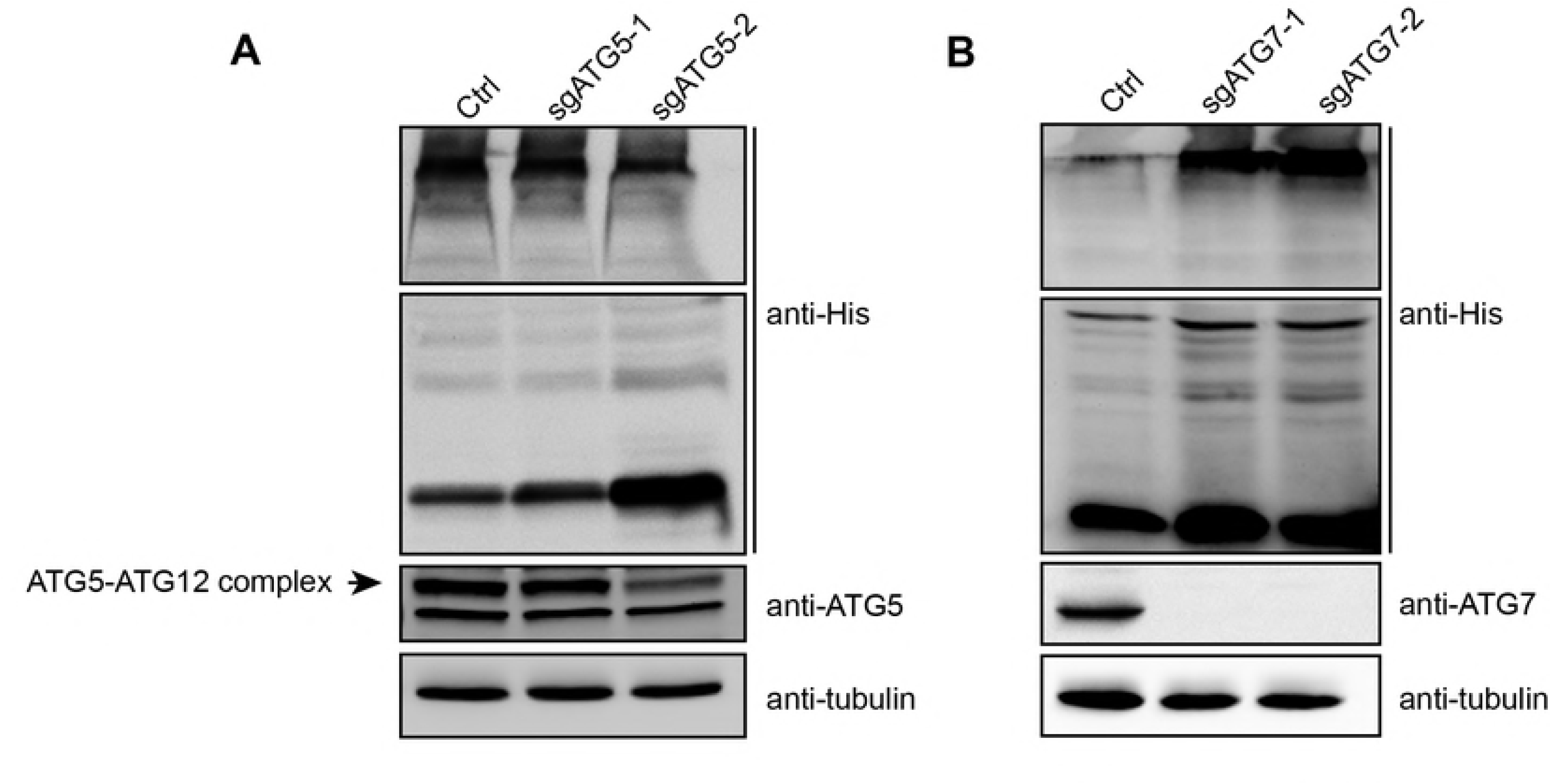
Suppressed expression of ATG5 and ATG7 leads to increased ASP expression. Two different 293T cell clones knocked out for either ATG5 (sgATG5–1 and sgATG5–2) (A) or ATG7 (sgATG7–1 and sgATG7–2) (B) and control stably transfected clones were transfected with an His-tagged NL4.3 ASP expression vector.. Cell lysates were prepared and analyzed by Western blot using anti-His, anti-ATG-5, anti-ATG7 and anti-tubulin antibodies. Monomeric and multimeric signals are representing different ASP signals. The signal for ATG5 is shown as the typical covalently linked ATG5-ATG12 complex.

The results thereby confirmed that ASP levels in transfected cells were modulated by autophagy, and that typical autophagy factors, i.e. ATG5-ATG12 and ATG7, were implicated.

**ASP interacts with autophagy factor p62**. Given that all tested ASP could induce autophagy, we next investigated on the possible mechanism of action. We first focused our attention on determining whether ASP could associate and co-localize with specific autophagy factors by confocal microscopy in different cell lines (Fig. 7). As we have previously reported (23), we first confirmed that expression of NL4.3 ASP in 293T cells showed association to LC3 upon co-immunoprecipitation with anti-ASP followed by Western blot analyses (Fig. 7A). In addition, ASP was confirmed to partially co-localize with LC3-II in both COS-7 and HeLa cells (Fig. 7B and data not shown). We next performed confocal microscopy experiment to identify potential co-localization with the important autophagy protein p62. Our results revealed that a clear co-localization between ASP and p62/SQSTM1 was observed in both HeLa and 293T cells (Fig. 7C).

**Figure 7.**
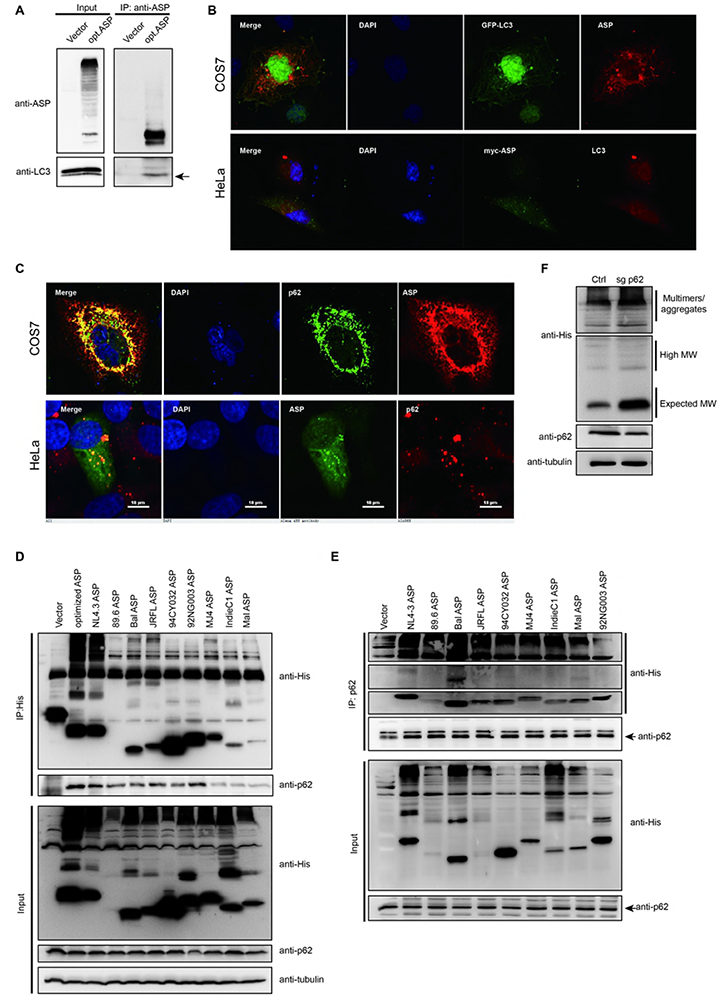
Association between ASP and p62 with and effect of knock out p62 expression on ASP levels. **A.** 293T cells were transfected with expression vectors for optimized NL4.3 ASP or the empty vector pcDNA3.1. After 48 h post transfection, immunoprecipitation was performed using the anti-ASP antibody and Western blot analyses were conducted through anti-ASP and anti-LC3 antibodies. The Mouse TrueBlot^®^ ULTRA (Anti-Mouse Ig HRP) (for ASP) was used as secondary antibody for Western blot analyses. **B.** Expression vectors pcDNA-opt-ASP and GFP-LC3 were transfected in COS-7 cells. After fixation and nuclei staining with DAPI, ASP was detected with anti-ASP antibodies followed by a goat anti-mouse IgG antibody coupled to Alexa Fluor 568, while LC3 detection was performed with an anti-LC3 antibody followed by a goat anti-rabbit IgG antibody coupled to Alexa Fluor 568. **C.** Expression vectors for NL4.3-ASP were transfected in COS-7 (upper panels) and HeLa (bottom panels) cells. Nuclei were stained with DAPI, and ASP was detected with anti-ASP antibodies followed by goat anti-mouse IgG antibodies coupled to Alexa Fluor 568 (COS-7) or Alexa Fluor 488 (HeLa), while p62 was detected by an anti-p62 antibodies followed by goat anti-rabbit IgG antibodies coupled to Alexa Fluor 488 (COS-7) or Alexa Fluor 568. **D-E.** Expression vectors of His-tagged ASP of various HIV clades were transfected in 293T cells and after 48 h, cellular extracts were used for immunoprecipitation with the anti-His (**D**) or anti p62 antibodies (**E**). Immunoprecipitated samples and total extracts were analyzed by Western blot using anti-p62, anti-His and anti-tubulin antibodies. **F.** 293T cell clone knocked out for p62 (sgp62) and control stably transfected clone were transfected with His-tagged NL4.3 ASP expression vector. Cell lysates were prepared and analyzed by Western blot using anti-His, anti-p62 and anti-tubulin antibodies. Monomeric and multimeric signals are representing different ASP signals.

Since ASP forms multimers and that p62 is an important cargo transporter of multimerized proteins toward the autophagy pathway, we next tested whether ASP and p62/SQSTM1 could indeed associate. 293T cells were transfected with expression vectors of His-tagged ASP of different clades and resulting extracts were used for co-immunoprecipitation with an anti-His antibody (Fig. 7D). Upon Western blot analyses, p62 indeed could co-immunoprecipitate with ASP in transfected 293T cells. To confirm these data, immunoprecipitation was performed in similarly transfected 293T cells with anti-p62 antibodies and analyzed with anti-His antibodies (Fig. 7E). These analyses confirmed the association between ASP and p62. The specificity of this interaction was further demonstrated in transfected 293T cells by the lack of interaction of ASP with the other autophagy factors, ATG5 and ATG7 implicated in the ASP-induced autophagy cascade (Fig. S3). The impact of this association between ASP and p62 was further tested through knockout of the p62 gene in 293T cells. Although two different targeted region were tested, only one strategy (sgp62-1) provided sufficient reduction in p62 abundance. Knockout cells were thus transfected with an ASP expression vector and analyzed by Western blot. These analyses indicated that reduction in p62 levels led to a marked increase in ASP abundance (Fig. 7F).

These data thus indicated that p62 co-localized and associated to the different ASP clade representatives, and that this association might consequently lead to induction of autophagy.

**Interaction domain of p62 and ASP ubiquitination**. We next sought to identify the domain of p62, which was responsible for its association with ASP. These domains have been previously shown to interaction with ubiquitinated protein, LC3 and p62 itself, and termed UBA, LIR and PB1, respectively (Fig. 8A). Various p62 mutants deleted for these various domains and fused to the GST protein were thus co-transfected with Myc-tagged NL4.3 or 94CY032 (clade A) ASP expression vector in 293T cells and resulting extracts were co-immunoprecipitated with anti-GST antibodies (Fig. 8B). Upon Western blot analysis of immunoprecipitated extracts, interaction of ASP (monomeric and multimeric) was shown to be importantly reduced in cells expressing the PB1 domain-deleted p62 mutant (Fig. 8B). Expression of other mutants (deleted for LIR or UBA regions) led to retention of ASP in extracts co-immunoprecipitated with anti-GST antibodies, thereby suggesting that these other domains were not crucial for the association between ASP and p62. Similar results were observed in immunoprecipitation experiments following co-transfection of the various p62 mutants and a His-tagged NL4.3 ASP expression vector in 293T cells upon Western blot analyses (Fig. S4).

**Figure 8.**
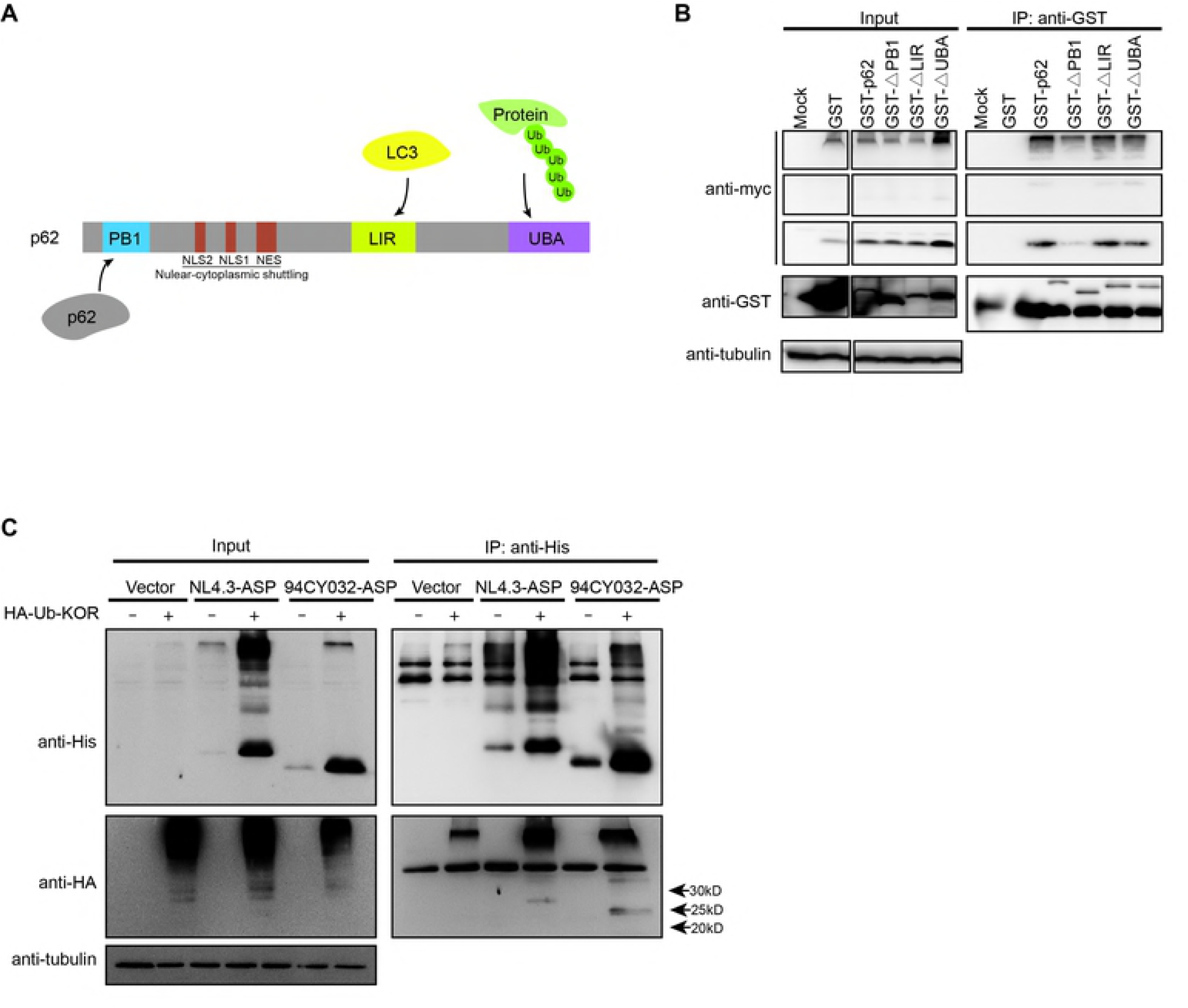
The interaction of p62 with ASP requires its homodimerization domain despite ASP ubiquitination. **A.** The various domains of the p62 protein are depicted and include the ubiquitination-binding domain UBA, the LC3-binding domain LIR, NLS and NES regions and the homodimerization domain PB1. **B.** 293T cells were co-transfected with expression vectors for GST-tagged wild-type and deletion mutants of p62 and of Myc-tagged-NL4.3 ASP. At 48 h post transfection, immunoprecipitation was performed using the anti-GST antibody and Western blot analyses of immunoprecipitated samples along with total extracts were conducted through anti-GST, anti-Myc and anti-tubulin antibodies. **C.** 293T cells were co-transfected with expression vectors for His-tagged NL4.3 or 94CY032 ASP and for the HA-tagged Ubiquitin-KOR mutant (restricted to monoubiquitination). At 48 h post transfection, immunoprecipitation was performed with anti-His antibodies and Western blot analyses of immunoprecipitated samples along with total extracts were conducted through anti-His, anti-HA and anti-tubulin antibodies. Arrows on the right side of the right panels indicated the position and expected MW (25–30 kDa) of the ubiquitinated ASP from the different tested proviral DNA.

Since p62/SQSTM1 has been often implicated in ubiquitination-dependent autophagy, it was surprising that the association between ASP and p62 did not involve the ubiquitinating interacting domain UBA. We were thus interested in determining if ASP was nonetheless ubiquitinated. Immunoprecipitation experiments were thus conducted in 293T cells co-transfected with two different His-tagged ASP expression vectors (from proviral DNA clade B NL4.3 and clade A 94CY032) and pRK5-HA-Ubiquitin-KOR (expressing an HA-tagged ubiquitin protein limiting targeted proteins to monoubiquitination). Upon transfection of 293T cells and immunoprecipitation with anti-His antibodies, Western blot analyses revealed that ASP was indeed ubiquitinated, as revealed by the expected size of a ubiquitinated monomeric ASP form in both clade A and clade B expressing cells (Fig. 8C). In addition, high molecular weight multimers were detected in immunoprecipitated extracts. Interestingly, for both tested ASP expression vectors, levels of the non-ubiquitinated form was also importantly increased in transfected cells expressing ubiquitin KOR, likely suggesting that these might represent non-ubiquitinated ASP protein composing ASP multimers targeted for ubiquitin-dependent degradation. In fact, high molecular signals were also importantly more abundant in KOR expressing cells. Further experiments using HA-tagged ubiquitin expression vectors selectively producing K48 (UPS-prone) or K63 (autophagy-prone) polyuquitination revealed that ASP was targeted by both of these modifications (data not shown).

Together, these data demonstrated that, despite the implication of the homodimerization domain of p62 in its association with ASP, ASP is ubiquitinated, which impacts its abundance in its monomeric and multimeric forms.

## Discussion

Antisense transcription in HIV −1 has been described and characterized in a number of studies (18). However, the encoded ASP protein remains a limited focus of investigation. We have formerly detected ASP in different cell types by various technical approaches and further demonstrated that this viral protein led to the induction of autophagy (7, 17, 23). In the current study, we demonstrate that ASP-associated autophagy is induced in all tested clade-derived ORF and is associated to p62 via its oligomerization domain. Furthermore, ASP was shown to be ubiquitinated, which impacted on its sensitivity toward degradation.

Based on our earlier studies linking ASP-induced autophagy to its multimerization capacity, we pursued our analysis of the ASP multimers and confirmed that ASP is indeed part of multimers by showing the interaction between two differently tagged ASP. As cysteine residues can mediate multimerization, denaturing agents were tested and indeed reduced the formation of the aggregates. Furthermore, removal of the cysteine-rich amino region of ASP led to similar lower abundance of detected high molecular signals with an increase in the monomeric version, especially noted in COS-7 cells. Interestingly, as expected, the amino-truncated version was also less prone to autophagy induction. It is not clear why increase in the abundance of amino-truncated ASP mutants was not as clearly seen in 293T cells, although higher selective degradation of the codon-optimized NL4.3 ASP monomers targeted by either degradative pathways in this cell type might account for these differences. However, differences in the levels of monomeric form of ASP deleted at their amino end was further supported by the lower abundance of multimers in cells expressing clade A ASP lacking the first 25 amino acid cysteine-rich region. These results argue that cysteine residues are implicated in the formation of these ASP multimers, which is reminiscent of other HIV-1 proteins (45, 46). Although our results argue that the cysteine triplet (position 10–12) are implicated in ASP multimer formation, we have not fully investigated on which cysteines are important but it is highly probable that various cysteine residues are involved, some of which could also lie outside of the tested regions. Furthermore, since autophagy was not completely abrogated in cells expressing amino deleted version of NL4.3 ASP, we speculate that other highly conserved regions (9) are likely to be involved in the multimerization of ASP.

In this study, a more representative assessment of the autophagy-inducing capacity was assessed through expression vectors of various ASP from various clade representatives of clinical isolates and laboratory strains. We first demonstrated for the first time that we can detect ASP from those different isolates at the expected molecular weight in transfected cells. Again, high molecular weight signals and multimers were prevalent in these analyses, albeit at different abundance, a variation potentially accountable by the absence of the cysteine-rich amino end in the case of clade A. Despite variation in multimer formation, all tested ASP induced autophagy, as determined by variation in the levels of p62 and LC3-II levels and by the presence of autophagosome-like signals by confocal microscopy. Although we have not tested all known clades (having limited out analyses to most frequent ones), our data argue for a conserved capacity of ASP from the different clade families to induce autophagy and that low abundance of multimers for certain ASP is nonetheless sufficient for this induction.

Our results thereby strongly suggest that ASP multimers in the form of potential aggregates induces autophagy and acts upon ASP abundance. This parallels similar aggregating proteins, known to be targeted for degradation by this pathway (47, 48). We have further confirmed the implication of autophagy on the abundance of ASP by targeting known autophagy factors through CRISPR-mediated deletion. Although certain selected clones still expressed varying levels of target protein, (likely reminiscent of either more than one clone and/or single allele inactivation), resulting ASP levels (monomer and multimers) were strongly affected and their increase in abundance inversely correlated with the remaining levels of the targeted protein. We have further looked at the potential mechanisms of autophagy induction by ASP and through localisation and immunoprecipitation experiments, an association between ASP and p62 was detected, being dependent on the PB1 homodimerization domain, but not affected in a p62 mutant deleted for its ubiquitin-interacting UBA region. Interestingly, this was comparable to previous studies, in which viral proteins could interact with p62 via a ubiquitin-independent manner (49, 50). In fact, HIV-1 Tat is one of these proteins and is selectively degraded through its association to p62 via autophagy in an ubiquitin-independent manner.

These latter results are unexpected given that other experiments have revealed that ASP is ubiquitinated and that such post-translational modification contributes to ASP degradation, as demonstrated by the use of the ubiquitin mutant preventing polyubiquitination. Expression of this polyubiquitination-blocking ubiquitin protein indeed led to an important increase in abundance of the non-ubiquitinated monomeric and multimeric forms of ASP, the former likely resulting from their its in multimer complex and subsequent release upon cell extract preparation. Hence, these multimers being composed of non-ubiquitinated and ubiquitinated ASP are thus subject to degradation by autophagy through their interaction with p62. Although it is not clear how ASP triggers autophagy, the recent publication of Lu *et al*. might shed light on the exact mechanism behind ASP-dependent induction of autophagy. The authors indeed argued that association of ubiquitinated multimeric proteins to p62 with subsequent induced autophagy is more dependent on the formation of aggregates than ubiquination itself (51). In this perspective, it is likely that the association of ASP with p62 depends on its oligomeric form and that the interaction with the ubiquitin-interacting UBA domain might not be detectable in our immunoprecipitation experiments as the association is more dependent on the multimeric form of ASP than the affinity toward ubiquitin conjugates. This is thus different from the previously suggested view that autophagy vs. UPS is mainly dependent on the type of linked ubiquitin in the polyubiquitination chain (K48 vs. K63) (52). Our results in fact argue that both types of modification are present in monomeric and multimeric forms of ASP (data not shown).

It is well known that autophagy is primarily responsible for the degradation of most long-lived proteins in cells, but also targets aggregated proteins as well as cellular organelles and infectious organisms (53). The importance of autophagy in modulating the abundance of ASP might be crucial to maintain its abundance at low level, as this protein is likely detrimental to cell survival (if in high excess), especially given its propensity to form important multimers. In addition, the capacity of ASP to function as an inducer of autophagy also depends on other concomitantly expressed viral proteins, which have been associated to this degradation pathway (39). As previously reported, Nef can inhibit autophagy at late steps of viral replication in macrophages and could thus block ASP-induced autophagy (5, 36). Another interesting aspect of our studies relate to the essential role played by autophagy in antigen presentation in cells, such as dendritic cells. Infected DC might thus be altered in their capacity to adequately process and present antigen, especially knowing that our previous study showed that ASP presented the highest expression in this cell type when compared to macrophages and activated CD4+ T cells (17).

Overall, our results demonstrate that the autophagy-inducing properties of ASP are observed with different clades ASP and that multimerization acts on autophagy induction. Our results further indicate that the association of p62 to ASP is likely essential for ASP and that ASP is ubiquitinated, also affecting the stability of the monomeric and multimeric forms of ASP. The body of results thereby helps to better appreciate the unstable nature of ASP in mammalian cells. The function of ASP in HIV-1 infection *in vivo* has yet to be defined but our results suggest that autophagy could link ASP to HIV-mediated pathogenesis and the chronic infection state. Future studies will be determinant in providing more information on these potential roles.

## Materials and Methods

### Plasmids

Expression vectors for the Myc-tagged optimized ASP and GFP-ASP optimized fusion proteins have been previously described (23). A series of deletion mutants (15 and 30 amino acids at amino or carboxyl ends) and internal deletion of cysteine triplet or PXXPXXP motif were generated by PCR from Myc-tagged optimized ASP expression vector using primers presented in Table S1. These resulted in the constructs termed pMyc-optimized-ASPΔN1-15, pMyc-optimized-ASPΔN1-30, pMyc-optimized-ASPΔC174-189, pMyc-optimized-ASPΔC159-189, pMyc-optimized-ASPΔ^10^CCC^12^ and pMyc-optimized-ASPΔPXXP. Proviral DNA p94CY032 (clade A), NL4.3 (clade B), JR-FL (clade B), 89.6 (clade B), MJ4 (clade C), p92NG003 (clade G) and p92NG083 (clade G) were obtained through the NIH AIDS Reagent Program (Germantown MD) while proviral DNA Mal (clade D) and IndieC1 (clade C) were provided by Dr. Anne Gatignol (McGill University, Montreal, Canada). DNA fragments encoding His-tagged antisense protein (ASP) were amplified by PCR from these proviral DNA or the Bal (clade B) envelope expression vector (provided by Dr. Michel J. Tremblay, Laval University, Quebec city, Canada) using specific forward and reverse primers bearing NotI and BamHI restriction site, respectively at their 5’ end. After NotI/ BamHI digestion, PCR products were inserted into the vector pTT5 (Youbio Inc. China: #VT2202,), which was similarly digested. CRISPR-Cas9 knockout plasmids (LentiCRISPRv2, Addgene: #98290 and psPAX2, AddGene: #12260) were constructed as previously described (54, 55). Pairs of oligonucleotides were designed for specificity toward targeted genes and are listed in Table S2. Annealing of each pair further led to the generation of BsmBI restriction sites, which were used to insert the resulting dimerized oligonucleotide in the BsmBI-digested LentiCRISPRv2 vector. Plasmids pDEST-GST-P62, pDEST-GST-P62-ΔLIR, pDEST-GST-P62-ΔUBA and pDEST-GST-P62-ΔPB1 were kindly provided by Dr. Lucile Espert (Université Montpellier, Montpellier, France) and express wild-type and various deletion mutants of p62 (56). The plasmids pRK5-HA-Ubiquitin-WT (Addgene: #17608), pRK5-HA-Ubiquitin-KOR, pRK5-HA-Ubiquitin-K48 and pRK5-HA-Ubiquitin-K63 and the empty vector (Addgene #17605) were gifts from Dr. Ted Dawson (Johns Hopkins University School of Medicine, Baltimore MD) (57). The pRK5-HA-Ubiquitin WT vector expresses HA-tagged wild-type ubiquitin, while pRK5-HA-Ubiquitin-KOR expresses a ubiquitin version in which all lysines were mutated to arginine. pRK5-HA-Ubiquitin-K48 and pRK5-HA-Ubiquitin-K63 encode for ubiquitin-mutated versions in which lysine 48 and 63 respectively have been preserved, the other lysine residues having been mutated to arginine.

### Cell lines and transfection

Human embryonic kidney 293T, African green monkey kidney COS-7 and human cervical HeLa cell lines were cultured in Dulbecco’s modified Eagle medium (DMEM) supplemented with 10% fetal bovine serum (FBS) (Wisent Bioproducts, St-Bruno, Canada). Cell lines were obtained from the American Type Culture Collection (ATCC) (Manassas VA). All DNA plasmids were transfected using Polyethylenimine (PEI) (Polysciences, Warrington PA) at a 7:1 ratio of PEI (μg): total DNA (μg) in FBS-free DMEM: empty vectors were added to normalize DNA quantity in between transfection and for transfection control samples. After an incubation of 20 min at room temperature, the PEI/DNA mixture was added to cells for 6 h, after which medium was removed and replaced by fresh supplemented DMEM medium. In certain transfection, cells were treated with Bafilomycin A1 (100 nM) for 6 h at 24 h post-transfection.

### Generation of CRISPR-Cas 9 stable cell lines

Pseudotyped lentiviruses were produced by co-transfecting 293T cells with lentiCRISPRv2 or lentiCRISPRv2-sgRNA (specific to ATG5, ATG7 or p62), the HIV-1 backbone-containing psPAX2 and the VSV envelope encoding-pVSVg vector (Addgene: #8454) using the PEI agent. At 36 h post-transfection, viral supernatants were filtered with 0.22um filters (Millipore Corporation, Billerica, MA) and added to 293T and COS-7 cells. CRISPR-Cas9 stable knockout cell lines were selected and maintained in DMEM supplemented with 10% FBS and 0.2ug/ml puromycin. Clones were generated by 10X serial dilution and single clone wells were further expanded in puromycin-containing medium. Selected and amplified clones were analyzed by PCR for the targeted region.

### Antibodies

Anti-GST antibodies were purchased from GE Healthcare Life Science (Chicago IL). Anti-Flag, anti-tubulin, anti-GAPDH, anti-LC3, anti-ATG5, anti-ATG7, anti-ATG12, anti-Beclin-1 and anti-p62 antibodies were purchased from Sigma-Aldrich (St-Louis MO). Antibodies against Myc, His and HA tags, HRP-conjugated anti-GFP and HRP-conjugated goat anti-mouse and anti-rabbit IgG antibodies were ordered from Santa Cruz Biotechnology Inc. (Dallas TX). Mouse TrueBlot^®^ ULTRA(Anti-Mouse Ig HRP) was purchased from Rockland Inc. (Limerick PA). Goat anti-mouse IgG coupled to Alexa Fluor 488 or to Alexa Fluor 568 were obtained from Thermo Fisher Scientific (Waltham MA). Monoclonal anti-ASP antibodies generated by Eurogentec (Liege, Belgium) were kindly provided by Dr. Jean-Michel Mesnard (Université Montpellier, Montpellier, France) and are specific to the amino acid sequence indicated in Fig. 4A.

### Western blot analyses and immunoprecipitation

Cells were washed in PBS 1X and then lysed in lysis buffer (Tris 50mM pH8.0, NaCl 100mM, EDTA 1mM, 1%Triton with proteinase inhibitor (Roche, Mississauga, Canada) on ice for 15 min. In certain experiments, β-mercaptoethanol (2 %) and varying concentrations of DTT (0.1 −0.5 M) were added to the lysis buffer. The lysates were centrifuged at 2000 × *g* for 15 min at 4°C. Proteins (the supernatant) were denatured by mixing 20 μl lysates with 4X loading solution (12% sodium dodecyl sulfate (SDS) with 3 mM Tris pH 6.8 and 0.05% bromophenol blue), followed by incubation at 70°C for 10 min. For immunoprecipitation, SureBeads™ Protein G Magnetic Beads (BioRad, Hercules CA) were resuspended (20 μl per sample) and washed twice with 500 μl cold 0.1% Tween20 in PBS (PBS-T). Antibodies diluted in 500 μl PBS-T (anti-Myc (1:200), anti-p62 (1:100), anti-His (1:200), and anti-GST (1:100)) were then added to the beads for 1 hour at room temperature, and beads were next washed three times with cold PBST. Total cell extracts (lysed in 50 mM Tris–HCl pH 8, 100 mM NaCl, 1 mM EDTA and 1% Triton) were then incubated with the antibody-bead complex overnight at 4°C. After three washing with cold PBS-T, bound fractions were eluted with 20 μl of 2X loading buffer heated at 100°C. Samples were run on 10–12% SDS-PAGE and blotted onto an Immun-Blot™ PVDF membrane (Bio-Rad Laboratories, USA) in PBS-T. Membranes were blocked with 5% milk in PBS-T at room temperature for 1 h and incubated with anti-ASP (1:2000), anti-GFP (1:5000), anti-Myc (1:250), anti-HA (1:10000), anti-GST (1:3000) anti-His (1:5000), anti-p62 (1:2000), anti-ATG5 (1:2000), anti-ATG7 (1:2000), anti-LC3 (1:500), anti-Beclin-1 (1:500), anti-GAPDH (1:1000) or anti-tubulin (1:10000), antibodies at 4°C overnight. Membranes were next incubated with the appropriate HRP-conjugated secondary antibodies (1 μg/ml) at room temperature for 2 h and visualized with the ECL Western blotting detection kit ((Immobilon™ Western, Millipore Corporation, Billerica MA). Image acquisition was performed with Fusion FX7 (Vilber Lourmat, Marne-la-Vallée, France). For certain experiments, densitometric analyses were conducted on LC3-related signals and a LC3-II/LC3-I ratio was calculated and normalized over cells transfected with an empty vector (set at a value of 1).

### Flow cytometry

Transfected COS-7 and 293T cells were washed with PBS, fixed with 4% formaldehyde for 10 min, and permeabilized with 0.1% Triton X-100 for 5 min at room temperature. Cells were then washed three times with PBS and incubated with the anti-Myc antibody (dilution, 1:250) overnight at 4°C. After three additional washes with PBS, cells were incubated with goat anti-mouse IgG antibodies coupled to Alexa Fluor 488 (1/500) (Invitrogen Canada Inc) for 1 h at 4°C. Cells were fixed with 1% formaldehyde and incubated overnight at 4°C before analysis with the FACScan device (BD Biosciences).

### Confocal microscopy

COS-7 and HeLa cells were seeded in 24-well plates containing a 1.5-mm-thick coverslip for 12 h and then transfected, as described above. At 48 h post-transfection, cells were rinsed twice with cold PBS and fixed with 4% formaldehyde/PBS for 10 min at room temperature on a slow rotatory shaker. Cells were next washed with PBS/ 0.1% Triton X-100 4 times. Fixed cells were incubated with a blocking solution (3% BSA, 5% milk and 50% FBS in 0.1% Triton X-100, 0.05% NaN_3_ PBS) overnight at 4°C, then rinsed with cold PBS three times and incubated with anti-p62 (1:250), anti-ATG5 (1:250), anti-ATG7 (1:250), anti-LC3 (1:250), anti-ASP (1:500), or anti-Myc (1:250) antibodies in a PBS solution containing 3% BSA, 0.1% TritonX-100 and 0.05% NaN_3_ at 4°C overnight. Cells were next washed with cold PBS/0.1% Triton X-100 and incubated with a secondary antibody (1:500 goat anti-mouse IgG coupled to Alexa Fluor 488 or goat anti-rabbit IgG coupled to Alexa Fluor 568) in the PBS solution described above for 45 min. at room temperature. Cells were washed with PBS/ 0.1% Triton X-100 and incubated in the immune-mount solution in the presence of 1% DAPI. All cell samples were visualized with a Nikon A1 laser scanning confocal microscope (Nikon Canada, Mississauga, Canada) through a 60X objective under oil immersion.

## Acknowledgments

We thank Denis Flipo for his excellent technical assistance.

---

**Table S1. Primers used for generation of ASP deletion mutants.** Primers are presented as forward and reverse pairs suitable for the generation of the deletion mutants by inverse PCR, as described in Material and Methods.

**Table S2: Primers for generation of expression vectors for CRISPR-mediated knockout of ATG5, ATG7, ATG12 and p62 genes.** Complementary oligonucleotides are depicted for each targeted gene and were inserted tin the lentiviral vector, as indicated in Material and Methods.

**Figure S1. Amino deletion and removal of a cysteine triplet in the amino end of ASP increase abundance of its monomeric form.** COS-7 cells were transfected with expression vectors for ASP mutants (see Fig. 2A and 3A) or the wild-type version vs. pcDNA3.1 (mock). At 48 h post-transfection, cellular extracts were analysed by Western Blot using anti-ASP and anti-tubulin antibodies.

**Figure S2. Expression of ASP from different clades leads to reduced levels of p62.** Expression vectors for His-tagged ASP from NL4.3, 94CY032, IndieC1 and Mal proviral DNA (vs. empty vector) were transfected in 293T cells and cell lysates were analyzed by Western blot for ASP (anti-His), p62 and tubulin. Clades of origin of the different tested ASP representative are indicated above each lane.

**Figure S3. Absence of association between ASP and the ATG5-ATG12 complex or ATG7.** The expression vector for His-tagged NL4.3 ASP (vs. empty vector) was transfected in 293T cells and at 48 h post-transfection, cellular extracts were used for immunoprecipitation with anti-His antibodies Immunoprecipitated samples and total extracts were analyzed by Western blot using, anti-His, anti-ATG5, anti-ATG7 and anti-tubulin antibodies.

**Figure S4. Interaction of p62 with ASP requires its homodimerization domain.** 293T cells were co-transfected with expression vectors for GST-tagged wild-type and deletion mutants of p62 and for His-tagged-NL4.3 ASP. At 48 h post transfection, immunoprecipitation was performed using the anti-GST antibody and Western blot analyses of immunoprecipitated samples along with total extracts were conducted through anti-GST, anti-His and anti-tubulin antibodies.

